# New tree shrew Parkinson’s model: a cost-effective alternative to monkey models

**DOI:** 10.1101/2023.09.01.555918

**Authors:** Hao Li, Leyi Mei, Xiupeng Nie, Liping Wu, Xiaofeng Ren, Longbao Lv, Jitong Yang, Haonan Cao, Jing Wu, Yuhua Zhang, Yingzhou Hu, Wenchao Wang, Christoph W. Turck, Bingyin Shi, Jiali Li, Lin Xu, Xintian Hu

## Abstract

The surge in demand for experimental monkeys has led to a rapid increase in their associated costs. Consequently, there is a growing need for the development of a cost-effective model for Parkinson’s disease (PD) that exhibits all core clinical and pathological phenotypes of PD. Evolutionarily, tree shrews (*Tupaia belangeri*) are much closer to primates in comparison to rodents and share more similar PD-related brain structures and movement ability with monkeys. As such, tree shrews represent an ideal small animal species for modeling PD. To develop a tree shrew PD model, we used the 1-Methyl-4-phenylpyridinium (MPP^+^) metabolite, derived from the well-established PD modeling drug 1-Methyl-4-phenyl-1,2,3,6-tetrahydropyridine (MPTP), to induce lesions in the dopaminergic neurons of the unilateral substantia nigra. After determining the optimal modeling dosage, the tree shrews consistently exhibited and maintained all classic clinical manifestations of PD for a 5-month period. The symptoms closely resembled the ones observed in PD monkeys and included bradykinesia, rest tremor, postural instability, and apomorphine-induced rotations, a classic phenotype of unilateral PD models. Immunostaining showed a significant loss of dopaminergic neurons (approximately 95%) in the substantia nigra on the lesioned side of the brain, a crucial pathological marker of PD. Further cytomorphological analysis revealed that the size of nigral dopaminergic neurons in tree shrews exceeded that of rodents and more closely approximated that of macaques. Based on the principle that structure determines function, the morphological similarity between tree shrews and monkeys may be an important structural basis for the manifestation of the highly similar phenotypes between monkey and tree shrew PD models. Collectively, this study successfully developed a PD model in a small animal species that faithfully recapitulated the classical clinical symptoms and key pathological indicators of PD monkeys. In addition to the well-recognized monkey models, the tree shrew model provides a novel avenue for the evaluation of PD treatments and underlying mechanisms.

## Introduction

Parkinson’s disease (PD) is a prevalent neurodegenerative disorder that affects approximately 2%–3% of the population aged 65 years and older (Poewe et al., 2017). The primary clinical symptoms of PD include bradykinesia, rest tremor, rigidity, and postural instability (Kalia and Lang, 2015). The characteristic feature of PD pathology involves the massive loss of dopaminergic neurons within the substantia nigra pars compacta. Furthermore, many individuals with PD also contain Lewy bodies, which mainly consist of α-synuclein aggregates, in the brainstem and cortex (Gibb et al., 1988; Spillantini et al., 1997). Currently, the precise etiology of PD remains elusive. Due to ethical regulations and associated risks, many essential experiments cannot be carried out on human subjects. Therefore, the development of animal models exhibiting stable and classical PD symptoms is crucial for effective PD treatment and mechanistic studies. To date, scientists have developed a variety of animal PD models, with rodent and non-human primate models being the most frequently used (Chen and Niu, 2019; Jiang and Dickson, 2018).

Due to their short lifespan, ease of maintenance, well-defined genetic background, and rapid reproductive cycle, rodents have been widely employed in the development of PD models, which has greatly contributed to PD research (Beal, 2001). However, rodents have limited fine and complex motor skills compared to humans. As a result, rodent PD models poorly replicate the core clinical symptoms of PD (Yang et al., 2021), including a total absence of rest tremor (Bobela et al., 2014). Motor dysfunctions in rodent PD models are typically not directly observable and are instead assessed through specially developed motor tests. For instance, the slowness of movement in 1-Methyl-4-phenyl-1,2,3,6-tetrahydropyridine (MPTP)-induced rodent PD models is commonly evaluated using the rotarod task (Kelly et al., 1998; Su et al., 2015), while the extent of nigral dopaminergic neuronal loss in 6-OHDA-induced hemi-parkinsonian rodent models is commonly reflected by apomorphine (APO)-induced rotational behavior (Bouchatta et al., 2018; Perlbarg et al., 2018).

In summary, given their evolutionary distance, notable distinctions exist between rodents and primates in terms of fine and complex motor behaviors. Rodents primarily exhibit limited movements across the two-dimensional plane at the base of their cages, while primates demonstrate a much broader range of three-dimensional motion and more complex movement skills. Moreover, significant variations exist in PD-related brain structures between these species. For instance, unlike primates, rodents lack the distinct separation of the caudate nucleus and putamen in the striatum, instead possessing an ensemble known as the caudate putamen (CPu) (Ni et al., 2018). These dissimilarities may contribute to the repeated failures of PD drugs developed and tested using rodent PD models in human clinical trials (Klivenyi and Vecsei, 2010). Therefore, although rodent models are essential for exploring the mechanisms underlying PD pathogenesis, they are not ideal for translational research aimed at bridging the gap between preclinical findings and clinical application in PD.

In contrast to rodents, monkeys represent the closest relatives to humans and share similar genetic backgrounds, brain structures, and behavioral phenotypes. In addition, we recently found that monkeys can spontaneously develop PD, providing a solid biological basis for their use as a PD model (Li et al., 2021a; Li et al., 2021c). Based upon this observation, we utilized the adeno-associated virus (AAV)-delivered CRISPR/Cas9 system to directly edit the *PINK1* and *DJ-1* genes in the unilateral substantia nigra of adult monkeys, thereby successfully establishing the first gene-edited PD monkey model recapitulating all classical phenotypes of PD. This breakthrough achievement has further validated the importance of monkeys as a model for studying PD (Li et al., 2021b). Consequently, the utilization of monkey models offers considerable advantages for clinical translation studies, as well as the exploration of PD etiology, identification of early diagnostic markers, and development of effective therapeutic interventions.

The well-established MPTP-induced PD monkey model not only reproduces the core clinical symptoms of human PD (Burns et al., 1983) but also exhibits massive dopaminergic neuronal loss in the substantia nigra, a core pathological feature of PD (Beal, 2001). Furthermore, our research team discovered that aged monkeys subjected to chronic MPTP administration also exhibit Lewy body-like pathology in the brain (Huang et al., 2018). Hence, the MPTP monkey model has been regarded as a very good PD model and widely used for PD treatment investigations. Nonetheless, this model has several limitations, including the instability of model symptoms and poor animal health.

Subsequently, we developed the first 1-Methyl-4-phenylpyridinium (MPP^+^)-induced PD monkey model, accomplished by direct injection of MPP^+^ into the substantia nigra. MPP^+^ is a metabolite of MPTP that exerts toxic effects by triggering oxidative stress through dopamine transporter reuptake, leading to dopaminergic neuronal death (Przedborski et al., 2000). This novel approach effectively addresses the limitations of the MPTP model (Lei et al., 2015). Using this model, we have successfully conducted stem cell transplantation and AAV-mediated PD gene therapy experiments that have yielded important insights. Despite all this, the use of macaque PD models is compromised by their lengthy growth and reproductive cycles, as well as the substantial expense associated with their acquisition and maintenance. Therefore, developing more cost-effective small animal models that faithfully replicate core PD phenotypes is of great importance and urgency.

The tree shrew belongs to the order *Scandentia* and is mainly found in Southeast Asia. It is an omnivorous animal with good climbing abilities and resembles a squirrel in appearance. Evolutionarily, tree shrews are the closest relatives to primates (Che et al., 2021; Fan et al., 2013; Fan et al., 2019; Savier et al., 2021; Ye et al., 2021). Several tree shrew models for depression (Wang et al., 2011), social avoidance (Ni et al., 2020), and virus infections (Kayesh et al., 2021; Xu et al., 2020; Yao, 2017) have been reported.

In comparison to rodents, tree shrews boast several advantages as candidates for PD modeling due to their evolutionary proximity to primates. Their foremost advantage lies in their complex, flexible, and three-dimensional spatial movement ability. Similar to macaques, tree shrews engage in extensive three-dimensional activities, including jumping and climbing, within a wide spatial range. Moreover, tree shrews exhibit remarkable motor skills (Fan et al., 2014), and exhibit a level of movement flexibility and complexity comparable to that observed in monkeys, thereby serving as a favorable foundation for developing PD animal models. Secondly, tree shrews possess a well-developed striatum with distinct separation of the putamen and caudate nucleus by the internal capsule, mirroring the striatal organization observed in monkeys (Ni et al., 2018; Rice et al., 2011). Moreover, the α-synuclein protein in tree shrews exhibits a striking similarity to human α-synuclein, showing a shared amino acid sequence of 97.1% (Wu et al., 2015). In contrast, rodents exhibit 95.3% similarity in amino acid sequence compared to human α-synuclein (Lavedan, 1998). Furthermore, the secondary structure and phosphorylation sites of α-synuclein in tree shrews are nearly identical to those in humans (Wu et al., 2015). Notably, the fibrillar form of α-synuclein in tree shrews can induce Ser129 phosphorylation in mouse neurons, similar to how human fibrillar α-synuclein behaves (Wu et al., 2019).

In contrast to monkeys, tree shrews have a smaller size, faster growth, and quicker reproduction. All this resulting in reduced maintenance costs and easier care requirements (Xiao et al., 2017). We submit that, for research purposes, the development of a tree shrew model of PD has great potential for substituting the use of monkeys (Yao, 2017).

In 2013, Ma et al. attempted to construct a tree shrew PD model via intraperitoneal administration of MPTP, representing the sole reported attempt to create such a model thus far. However, the clinical symptoms observed in these tree shrews were not persuasive due to the absence of quantitative analysis of PD-related behaviors and lack of classical PD pathology, such as significant loss of nigral dopaminergic neurons (Ma et al., 2013). Thus, the model did not present a compelling representation of PD. Moreover, none of the animals under investigation survived beyond five days subsequent to MPTP administration, which could be attributed to the administering technique involving the systemic introduction of highly toxic MPTP.

In the present study, we leveraged our experience in developing a hemi-parkinsonian model of MPP^+^-induced PD in rhesus monkeys (Lei et al., 2015) to establish a tree shrew PD model. We first determined the appropriate dose of MPP^+^ required for the model, followed by the precise injection of the optimal dose into the unilateral substantia nigra of the tree shrews. This modeling strategy has several advantages. Firstly, the precise administration of MPP^+^ directly into the substantia nigra facilitated a targeted and well-defined range of damage, minimizing non-specific harm and ensuring the overall well-being of the model animals. Secondly, this approach led to the development of a self-controlled hemi-parkinsonian model, thereby reducing the number of required animals, and minimizing individual variations.

## Methods

### Animals and ethics

Prior to conducting this study, ethical approval was obtained from the Laboratory Animal Welfare and Ethics Committee of Kunming Institute of Zoology, Chinese Academy of Sciences (SMKX-20190106). Eleven male tree shrews (Table 1) were individually housed in stainless steel cages (395 × 300 × 595 mm) containing a nest box (246 × 158 × 147 mm) in a feeding room maintained at a temperature of 25–27 °C, humidity of 55%–70%, and controlled lighting conditions of 12 h dark/12 h light (dark: 20:00–08:00; light: 08:00–20:00) (Chen et al., 2022; Meng et al., 2016). The tree shrews were provided with adequate food and free access to water under the care of experienced veterinarians. The selection criteria for inclusion in the study were as follows: 1) male; 2) approximately 1.5 years old; 3) in good health, with a weight range of approximately 130–150 g; and 4) no prior experimental experience or abnormal behavior. Trained caretakers gently handled the tree shrews prior to any procedures, such as drug administration and anesthesia, to mitigate stress levels and promote animal well-being.

**Table 1.** Detailed information on 11 tree shrews used in the experiments.

| Animal | Experimental purpose | MPP<sup>+</sup> dose (μg) | Weight (g) | Sex | Age (years) |
| --- | --- | --- | --- | --- | --- |
| TS 1 | Dose exploration | 5 | 140 | Male | 1.5 |
| TS 2 | Dose exploration | 36 | 130 | Male | 1.5 |
| TS 3 | Dose exploration | 60 | 140 | Male | 1.5 |
| TS 4 | Dose exploration | 60 | 141 | Male | 1.5 |
| TS 5 | Dose exploration | 80 | 130 | Male | 1.5 |
| TS 6 | Dose exploration & modeling | 50 | 133 | Male | 1.5 |
| TS 7 | Dose exploration & modeling | 50 | 150 | Male | 1.5 |
| TS 8 | Modeling | 50 | 146 | Male | 1.5 |
| TS 9 | Modeling | 50 | 146 | Male | 1.5 |
| TS 10 | Modeling | 50 | 137 | Male | 1.5 |
| TS 11 | Modeling | 50 | 146 | Male | 1.5 |

### Experimental design

The experiment was conducted in two stages. In Stage 1, seven tree shrews (TS 1–TS 7) were used to determine the appropriate dose of MPP^+^ (MPP^+^ iodide, D048, Sigma, USA) required for PD model development. This dose was based on the survival of tree shrews and stability of PD-related clinical symptoms. Ultimately, a single injection of 50 μg of MPP^+^ was deemed as a suitable modeling dose. In Stage 2, four additional tree shrews (TS 8–TS 11) were added to the study cohort, and the formal modeling procedure was conducted (Figure 1). Experimental design details are presented below.

**Figure 1:**
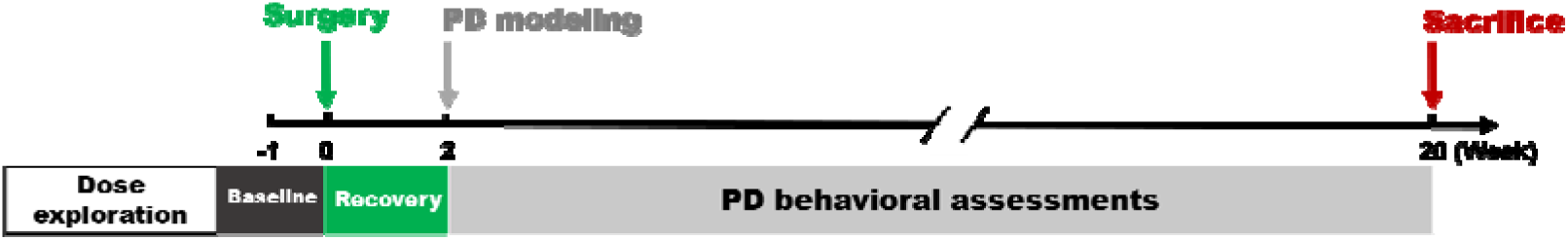
Experimental procedure. During modeling, baseline tree shrew data (black box) were collected one week before the operation, followed by a 2-week recovery period (green box) post operation. Behavioral data collection (gray box) spanned postoperative weeks 2 to 20. After 20 weeks, the tree shrews were sacrificed for pathological tests. The dose exploration experiments were consistent with the formal modeling of PD, except behavioral data collection lasted only four weeks.

#### Stage 1: Determining appropriate dosage of MPP^+^ for injection

As this was the first attempt to develop an MPP^+^-induced PD model in tree shrews, it was crucial to establish the appropriate dosage prior to commencing formal modeling procedures. Seven male tree shrews (TS 1–TS 7) were used in this experiment. In general, excessive doses of MPP^+^ lead to animal death, while low doses lead to significant recovery of PD symptoms. Considering the average mass of a monkey brain (100 g) in relation to that of a tree shrew (3 g) and referring to the dose of MPP^+^ used in our previous study on monkey brains (100 mg/mL–8 μL) (Lei et al., 2015), we estimated that the appropriate MPP^+^ dose for tree shrews was 24 μg. The minimum and maximum doses were subsequently determined within a range three times below or above the estimated dosage, resulting in a range of 8–72 μg. Therefore, five doses were used in this experiment, including 40 mg/mL–2 μL (80 μg), 30 mg/mL–2 μL (60 μg), 25 mg/mL–2 μL (50 μg), 20 mg/mL–1.8 μL (36 μg), and 10 mg/mL–0.5 μL (5 μg).

Determination of the appropriate dosage relied on two key indicators: i.e., survival probability, referring to the percentage of tree shrews that survived following administration of different doses of the drug, and appearance of classical PD symptoms and their stability. Evaluation was conducted through weekly video recordings during the initial month after MPP^+^ injection, followed by recordings once every two weeks. The PD symptoms exhibited by tree shrews were scored using the improved *Kurlan* scale (part A), a widely recognized scale for evaluating parkinsonian symptoms in monkeys.

Currently, there is no specific scale available for evaluating motor symptoms in tree shrews with PD. However, considering the similarity in the preference for broader three-dimensional play spaces observed in both tree shrews and monkeys, as well as the high mobility and comparable levels of flexibility and complexity demonstrated by tree shrews, as indicated in previous study (Fan et al., 2014), we used the improved *Kurlan* scale (part A) to quantify PD motor symptoms in tree shrews, which yielded excellent outcomes.

#### Stage 2: Tree shrew PD modeling

After establishing the optimal dosage (unilateral nigral injection of 50 μg MPP^+^), four additional tree shrews (TS 8–TS 11) were used to develop a hemi-parkinsonian model. The modeling period was the same as that of the two PD tree shrews (TS 6–TS 7) successfully induced using the same dosage in Stage 1. Throughout the modeling process, behavioral videos of the tree shrews and APO-induced rotation data were collected every two weeks.

To quantify the clinical symptoms of PD and APO-induced rotation counts, the behavioral video recordings were analyzed by two trained observers blind to the experimental conditions. In addition, brain tissue samples from the tree shrews were obtained at week 20 of modeling for pathological staining and quantification. The resulting data were compared with those derived from macaques and rodents. Details on the technical methods involved are described below.

### Experimental methods and techniques

This section comprises the following components: MPP^+^ injection into the unilateral substantia nigra, quantification of PD symptoms, pharmacological validation through APO-induced rotation, PD pathological study, data analysis, and comparison of morphological parameters of dopaminergic neurons from different species.

#### Stereotactic injection of MPP^+^ into substantia nigra

The surgical procedures performed on the tree shrews followed the experimental guidelines for neurotoxin-induced PD animal models (Tieu, 2011), as well as our previous experiments. Prior to the operation, each tree shrew was gently removed from its cage by the experimenter, and its hindlimb was intramuscularly injected with pentobarbital (80 mg/kg, 40 mg/mL). Subsequently, the head was secured on a stereotaxic instrument (68509, stereotactic instrument for rats with double tracks and double arms, RWD, China) under anesthesia (isoflurane mixed with pure oxygen (2% concentration)).

After each tree shrew was anesthetized, its head was shaved, and the scalp was disinfected using iodophors and alcohol. Subsequently, a careful incision was made on the scalp to expose the underlying skull, which was then disinfected using 3% hydrogen peroxide (H_2_O_2_) followed by a thorough saline wash. The precise coordinates for targeting the right substantia nigra pars compacta (AP = −3.88 mm, ML = 2.00 mm, DV = 10.00 mm) were determined according to a previously published tree shrew brain atlas (Jiang-Ning Zhou, 2016).

Before administering MPP^+^, the precise location of the substantia nigra was identified for each tree shrew by referencing the bregma and calculating the coordinates for inserting the injection into their skull. Subsequently, a hole was drilled in each skull, and a glass microtube (World Precision Instruments Inc., WPI, USA) loaded with the MPP^+^ solution was positioned accordingly. The microtube was then slowly lowered into the brain by manipulating an arm of the stereotaxic instrument, ensuring adherence to the depth specified in the tree shrew brain atlas.

Using a microsyringe pump, the MPP^+^ solution was injected into the right substantia nigra pars compacta of the tree shrew via the glass microtube at a rate of 100 nL/min. After the injection, the microtube was kept stationary for 5 min before being slowly withdrawn. A self-control was established by injecting an equivalent volume of saline into the left substantia nigra pars compacta of the tree shrew using the same procedure. Upon completion, the skull was rinsed with saline, and the scalp was sutured and disinfected with iodophor. Following surgery, the tree shrews were placed in a warm environment and returned to their nest box upon waking.

#### Quantification and analysis of PD clinical symptoms

The appearance of core PD symptoms serves as a critical indicator for evaluating the successful establishment of an animal model. For this purpose, a digital video camera (SONY HDR-XR260, Japan) was used to record daily free movement of the tree shrews in their cages. These video recordings, lasting 1 h each from 09:00–10:00 am, were analyzed by two trained observers. Baseline behavior was recorded one week before surgery, and subsequent recordings were conducted every one to two weeks, as needed, for five months. The dose exploration experiment was conducted over a period of one month.

The recorded videos were analyzed using the improved *Kurlan* scale (part A). This scale consists of seven items with corresponding scoring criteria: i.e., Tremor (0–3), Posture (0–2), Gait (0–4), Bradykinesia (0–4), Balance (0–2), Gross motor skills (0–3), and Defensive reaction (0–2). Each item is scored according to the severity of the specific symptom, and the total PD score is obtained by summing the scores of all seven items. The maximum possible total PD score is 20, with a higher score indicating more severe motor symptoms associated with PD.

The motor symptoms exhibited by the tree shrews were evaluated based on the seven items listed above. Two independent observers, blind to the experimental conditions, simultaneously scored each video. In cases of observer discrepancies, an experienced experimenter was consulted to review the videos, discuss the observations, and reach a consensus before scoring. Due to the temporary suspension of laboratory animal-related operations because of the COVID-19 pandemic, behavioral video recordings of the four tree shrews (TS 6–TS 7) in Stage 2 were only collected at baseline, and weeks 2, 4, and 20 following the operation. Despite this interruption, it did not impact the validity of our conclusions, as detailed below.

#### APO-induced rotation counts in tree shrews

APO, a commonly used drug for evaluating behavioral pharmacological performance in animal models of hemi-parkinsonism (Djamshidian and Poewe, 2016), was used in this experiment. Initially, tree shrew behaviors were video recorded for 1 h (09:00–10:00 am) while they freely moved within their home cages. Subsequently, the tree shrews were carefully removed by an experimenter and administered an intramuscular injection of APO (1 mg/mL, 0.2 mL, 013-18323, Wako, Japan) in their hindlimb. After the injection, the tree shrews were returned to their respective cages and observed for an additional hour (from 10:05–11:05 am). Finally, the number of APO-induced rotations exhibited by each tree shrew was counted.

An APO-induced rotation was defined as the completion of a full circle (360°) in either a clockwise or counterclockwise direction, with the initial direction arbitrarily determined before counting. The tree shrews were then manually rotated by 360° in either a clockwise or counterclockwise direction, which was scored as +1 rotation in the respective direction. For the two tree shrews (TS 6–TS 7) involved in the dose exploration and modeling stages, data collection was completed for all time points. However, due to the COVID-19 pandemic, data for the four tree shrews (TS 8–TS 11) introduced at Stage 2 only included rotation data collected at baseline and weeks 2, 4, and 20.

#### PD pathology tests

##### a) Brain tissue fixation and vibratome sectioning

At week 20 (five months) of PD modeling, the six tree shrews (TS 6–TS 11) that underwent the modeling process were euthanized using an intramuscular overdose of pentobarbital sodium (100 mg/kg, 40 mg/mL). The tree shrews were perfused with 0.01 M phosphate-buffered saline (PBS) and 4% paraformaldehyde via the heart to fix their brains, which were subsequently extracted and post-fixed in 4% paraformaldehyde for 48 h. Given that the substantia nigra is the most important brain region for pathological changes in PD, the tree shrew brain atlas (Jiang-Ning Zhou, 2016) was used to determine its specific location, i.e., anterior to posterior (AP) extension from bregma −3.75 mm to bregma −5.89 mm and total AP length of 2.14 mm. Tissue blocks of the substantia nigra were then embedded in 3% agarose and sectioned using a vibratome (Leica, VT1000S, Germany).

To ensure comprehensive coverage of the substantia nigra pars compacta, the cutting block for sectioning was adjusted to an AP length of 2.2 mm, with each section being 40 μm thick. The aim was to obtain approximately 55 brain sections from each tree shrew. However, due to individual variations and instrument inaccuracies, approximately 50 coronal sections of the substantia nigra were obtained from each tree shrew. To create a complete set of coronal sections of the substantia nigra, five consecutive brain sections from each tree shrew were collected in the same Eppendorf tube (1.5 mL) containing tissue antifreeze solution. This resulted in approximately 10–11 tubes, which were stored in a −20 °C freezer for further use.

To compare the staining results of our experiment with those of monkeys, substantia nigra sections from six macaques with unilateral substantia nigra MPP^+^ lesions from another experiment (IACUC19004) were used. The monkey brain tissue was fixed using 4% paraformaldehyde for approximately four weeks. Subsequently, tissue blocks of the mesencephalic substantia nigra were embedded using 3% agarose and sectioned using a Leica VT1000S vibratome. According to “The Rhesus Monkey Brain in Stereotaxic Coordinates” (Paxinos, G., Huang, X. & Toga, A. W. 1999, San Diego, USA: Academic Press.), the substantia nigra pars compacta in macaques extends from bregma −9.90 mm to bregma −17.78 mm, with an AP length of 7.88 mm. To determine the appropriate section blocks for the macaque substantia nigra, 3T magnetic resonance imaging (MRI) scans were conducted on monkey brains using a uMR790 MRI system (United Imaging, Shanghai, China) at the Kunming Institute of Zoology, Chinese Academy of Sciences, which revealed an AP length range of 7.0–7.9 mm. Thus, to ensure full coverage of the macaque substantia nigra, an AP length of 8.0 mm was selected for the section blocks. Each slice was 40 μm thick, and it was anticipated that 200 brain sections would be obtained for each macaque. However, due to individual variations in macaque specimens and instrument inaccuracies, approximately 160–180 midbrain substantia nigra coronal sections were obtained per macaque, forming a complete set of substantia nigra coronal sections for each macaque. Ten consecutive brain sections per macaque were collected in the same Eppendorf tube (1.5 mL) containing tissue antifreeze solution. Overall, 16–18 tubes per macaque were collected and stored in a −20 °C freezer for further analysis.

##### b) Immunohistochemical staining

To obtain representative sections of the midbrain substantia nigra from the tree shrews, we ensured that the selected brain slices covered the entire AP substantia nigra area. One brain slice was selected from each of the 10 centrifuge tubes containing preserved substantia nigra sections. This resulted in a total of 10 brain slices for tyrosine hydroxylase (TH) immunohistochemical staining, providing 60 brain slices from the six tree shrews. As a self-control, the same section from the other lateral substantia nigra was used. Similarly, we selected 10 brain slices per macaque for TH immunohistochemical staining.

The TH immunohistochemical staining protocol followed previous research (Li et al., 2015). The sections were treated with 3% H_2_O_2_ (5 min, Maixin, Fuzhou, China), 3% triton X-100 (4 min, Solarbio, Beijing, China), and 10% goat serum (15 min, Maixin, Fuzhou, China). The sections were then incubated with a rabbit anti-TH antibody (1:1LJ000, AB152, Millipore, USA) at 4 °C overnight, then with an anti-rabbit/mouse secondary antibody (PV-9000 kit, ZSGB-BIO, Beijing, China) the following day for 30 min at 37 °C. After this, the sections were incubated with 3,3’-diaminobenzidine (DAB 1031 kit, 20×, Maixin, Fuzhou, China) for 2 min and counterstained with hematoxylin for 1 min.

Following the staining process, the sections were subjected to ethanol gradient dehydration, xylene transparency, neutral resin blocking, and air-drying. Stained images of each section were then collected using a digital pathology section scanning system (SQS-40P, Shenzhen Shengqiang Technology Co., Ltd., China). The system allowed for image acquisition at different magnifications: 1× (13.62 mm × 8.95 mm), 4× (3418.90 μm × 2246.64 μm), and 20× (804.38 μm × 442.50 μm), enabling visualization of the substantia nigra region. Additionally, at a higher magnification of 40× (402.50 μm × 221.25 μm), images of the substantia nigra regions on both the MPP^+^-lesioned and healthy sides were captured for cell counting. TH-positive neurons in the substantia nigra on the healthy and MPP^+^-lesioned sides were labeled using ImageJ software (National Institutes of Health, Bethesda, Maryland, USA). The TH-positive cells with clear nuclei and intact soma were counted.

#### Morphological parameters of dopaminergic neurons in tree shrews

The average diameter of the soma and nucleus of dopaminergic neurons in tree shrews was measured using ImageViewer (Shengqiang Technology Co., Ltd., Shenzhen, China). Cells in the substantia nigra of the healthy side were selected from 60 brain slices of the tree shrews. Specifically, one morphologically intact cell was chosen from each brain slice, located near the center of the long axis of the substantia nigra, to represent the population. This resulted in a total of 60 analyzed cells.

Measurement of soma diameter involved identifying dopaminergic neurons in the substantia nigra pars compacta that contained 2–3 neurites and had spindle-shaped soma. The mean values of the long and short axes of the soma were measured, both through the nucleus and perpendicular to each other. Specifically, following the protocol described by Nelson et al. (Nelson et al., 1996), the mean diameter of the soma was calculated as ½ of the sum of the long axis and short axis measurements.

For nucleus diameter measurement, most cell nuclei are round (Webster et al., 2009) and nucleus diameter measurement through the nuclear and nucleolar plane is simple and adequate (Scaravilli and Jacobs, 1981). Therefore, straight distance through the nucleolus was used as the diameter of individual nuclei. The same measurement method was applied to determine mean diameter of the soma and nucleus of dopaminergic neurons in the substantia nigra pars compacta on the healthy side of the macaques.

The same methods described above were used to obtain the morphological parameters of dopaminergic neurons in the substantia nigra of C57 mice, including average diameter of the soma and nucleus. Clear images of TH immunohistochemically or immunofluorescently stained dopaminergic neurons in the substantia nigra of C57 mice were obtained from previous research (Gonzalez-Cabrera et al., 2017; Lin et al., 2020; Tiklova et al., 2019; Won et al., 1989). Similarly, using the same methods, the average diameter of the soma of dopaminergic neurons in the substantia nigra of Wistar rats was obtained from relevant literature (Bigham et al., 2021; De March et al., 2006; Guan et al., 2000; Karunasinghe et al., 2017; Van der Perren et al., 2011).

### Statistical analysis

Data analysis was conducted using SPSS v25 and GraphPad Prism v8.0. Analysis of variance (ANOVA) was used to compare total PD scores of the six tree shrews (TS 6–TS 11) at different modeling times. The nonparametric Mann-Whitney U test was used to compare the scores of each of the seven motor symptoms between the baseline and modeling stages at weeks 2 and 20 in the six tree shrews. Similarly, the Mann-Whitney U test was utilized to examine differences in the number of APO-induced rotations at baseline and weeks 2 and 20 of modeling among the six tree shrews (TS 6–TS 11). Additionally, the Mann-Whitney U test was used to determine differences in TH-positive cell density in the bilateral substantia nigra of the six tree shrews (TS 6–TS 11). The *t*-test was used to analyze differences in the mean diameter of the soma and nucleus of nigral dopaminergic neurons (n = 60) between C57 mice, Wistar rats, tree shrews, and macaques.

## Results

### Optimal MPP^+^ modeling dosage

During the dosage exploration experiments, we administered five different doses of MPP^+^ into the right substantia nigra of tree shrews, based on previous macaque models. Results showed that the tree shrews were able to tolerate MPP^+^ doses of 5 μg, 36 μg, and 50 μg, but rapidly died when the dose exceeded 50 μg (TS 3–5, Figure 2A). However, the 5 μg and 36 μg doses did not induce stable PD clinical symptoms, with PD scores decreasing to 1–2 at week 4 of modeling (see Methods). In contrast, the two tree shrews injected with the 50 μg dose showed stable PD clinical symptoms, with scores of 8–9 (Figure 2B). Therefore, we concluded that the appropriate dose for establishing a tree shrew PD model by unilateral nigral injection of MPP^+^ was 50 μg.

**Figure 2:**
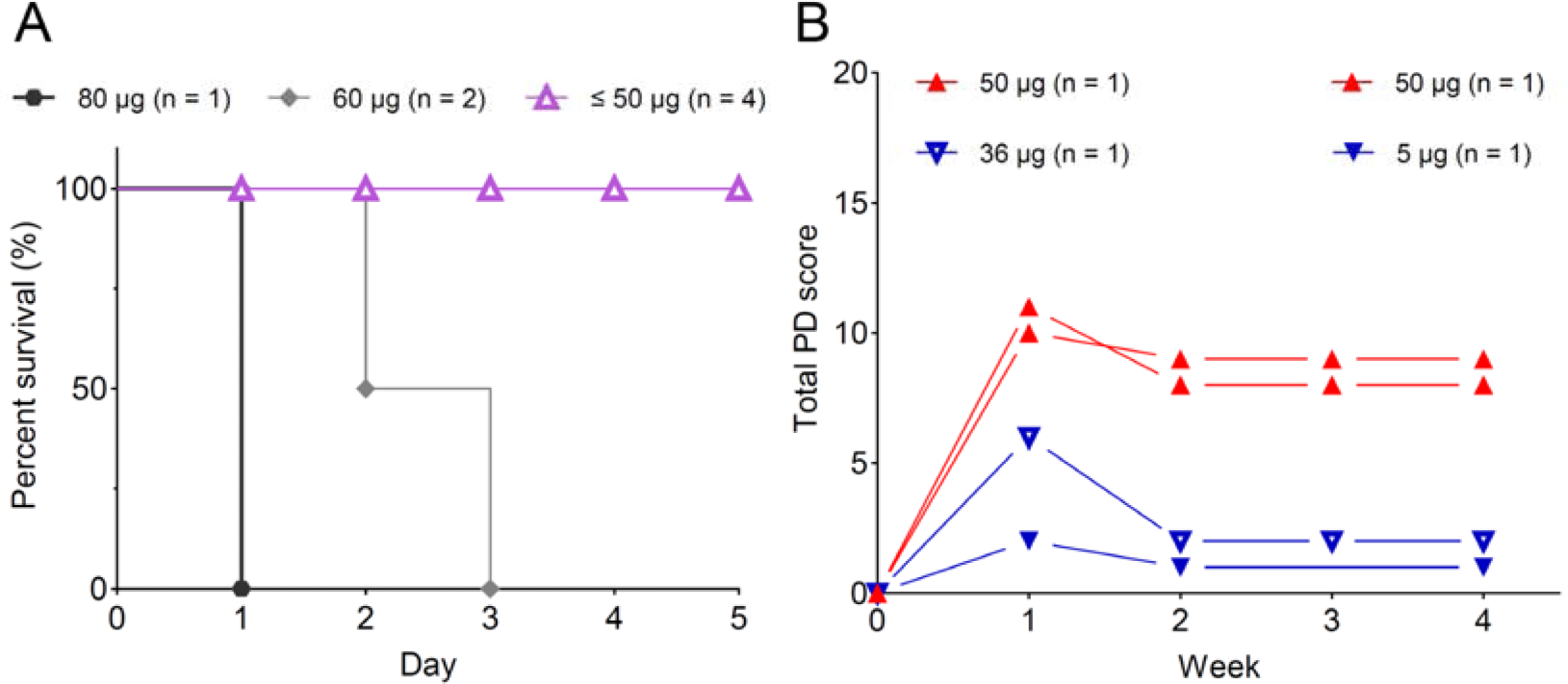
Exploration of appropriate dose for developing a tree shrew PD model based on MPP^+^ injection into the unilateral substantia nigra (n = 7). (A) Relationship between MPP^+^ dose and mortality. At an MPP^+^ dose of 80 μg, the tree shrew died within one day of surgery. At 60 μg, two tree shrews died three days after surgery. All surviving tree shrews received doses of 5 μg, 36 μg, or 50 μg, indicating that the appropriate dose for developing a tree shrew PD model by MPP^+^ injection into the unilateral substantia nigra ranged from 5–50 μg. (B) Relationship between MPP dose and PD score stability. One week after surgery, the PD scores increased in all tree shrews receiving doses of 5 μg, 36 μg, and 50 μg, reaching 2, 6, and 10–11, respectively. At four weeks post-surgery, the scores decreased to 1, 2, and 8–9, respectively, and remained stable and consistent until week 20 (see Figure 4 for details). Therefore, the appropriate dose for developing a tree shrew PD model by unilateral nigral injection of MPP^+^ was determined to be 50 μg. Note: In Figure B, behavioral video recordings were not collected at postoperative week 3 for the tree shrew receiving the 5 μg dose due to a cage change, which may disturb behavior.

### Formal tree shrew PD modeling

Based on a 50-μg modeling dose, we formally modeled PD in four additional tree shrews (TS 8–TS 11) over 20 weeks. As the modeling parameters of these four tree shrews were the same as those of the two tree shrews (TS 6–TS 7) in the previous dose exploration experiment, we combined the modeling data of these six tree shrews.

### Manifestation of PD clinical symptoms in tree shrew modeling

Upon completion of the PD model, we conducted an initial evaluation of the clinical symptoms of PD in the six tree shrews receiving the appropriate modeling dose (TS 6–TS 11). Motor symptoms are crucial diagnostic indicators for PD patients and for determining the successful development of animal PD models (Sedelis et al., 2001; Sveinbjornsdottir, 2016). For example, compared to its normal condition (Figure 3A, Video S1), TS 11 displayed several motor deficits, including difficulties in forelimb use (Figure 3B), balance issues (Figure 3C), and stooped posture (Figure 3D), two weeks after the induction of MPP^+^ lesions in the right substantia nigra.

**Figure 3:**
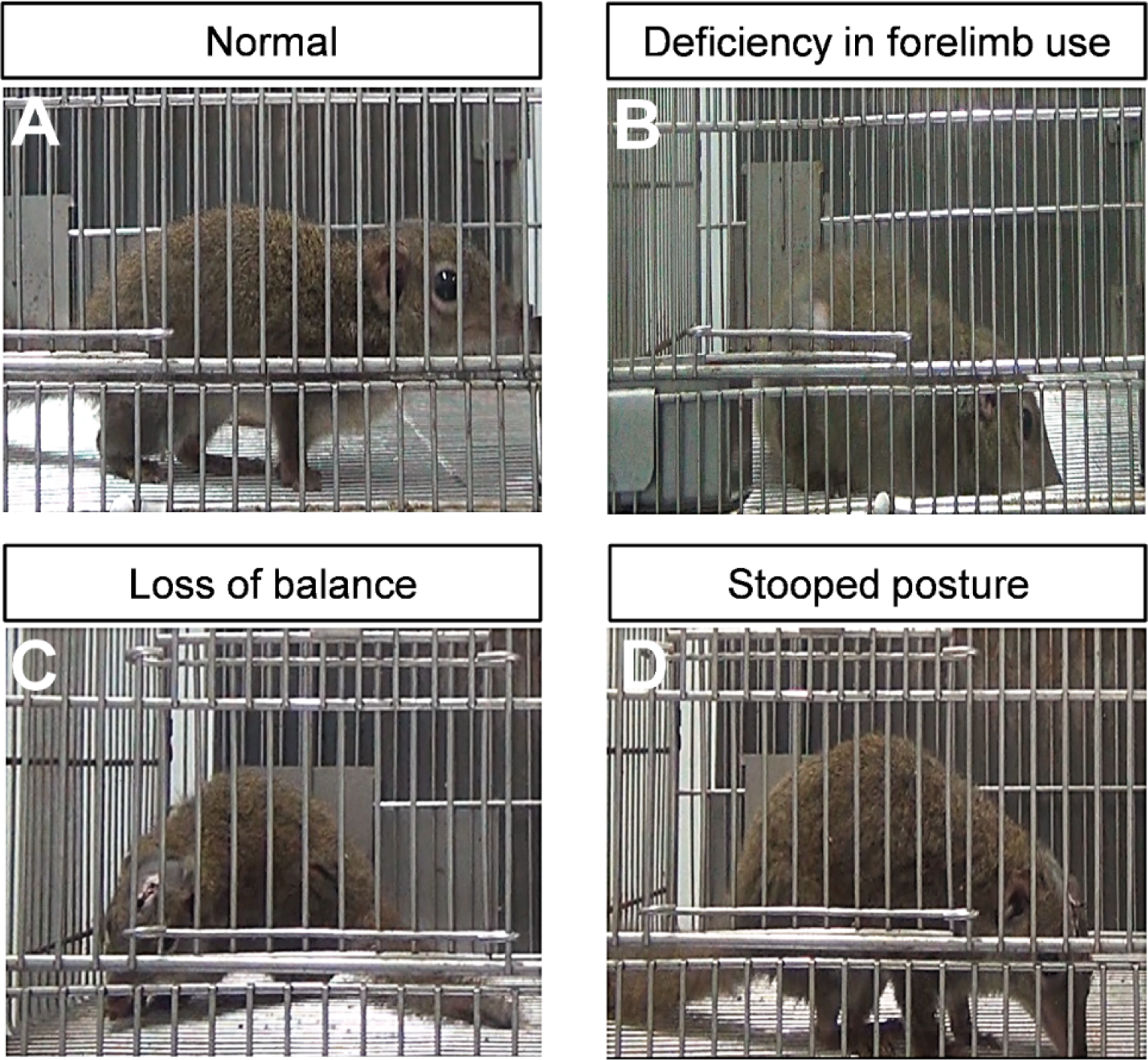
Significant PD-characteristic motor deficits in TS 11 at two weeks after a single 50-μg dose injection of MPP^+^ into the right substantia nigra. (A) Normal posture. (B) Difficulty in upper limb use, with frequent falling. (C) Balance issues, with falling to the left side. (D) Hunched and dorsally flexed posture, with spine uplifted and face adherent to the ground.

We observed distinct clinical motor symptoms of PD, including bradykinesia (Video S2), rest tremor (Video S3), and gait abnormalities, in the recorded videos. These symptoms align with the diagnostic criteria for PD established by the UK Parkinson’s Disease Society Brain Bank, which states that the presence of bradykinesia, along with at least one of the following symptoms - rigidity, rest tremor, or postural instability - is indicative of PD (Kalia and Lang, 2015; Moustafa et al., 2016; Poewe et al., 2017; Tolosa et al., 2006). Thus, the tree shrew PD model established by MPP^+^-induced lesions in the unilateral substantia nigra faithfully meets the clinical diagnostic criteria set for human PD.

Similar to macaques, tree shrews show a preference for a wide range of three-dimensional space and are recognized for their distinctive motor performances, particularly jumping and quick climbing. However, in our study, these habitual movements were no longer observed in the tree shrews after two weeks of PD modeling (Video S4). This absence suggests a marked reduction in locomotor speed and amplitude, effectively mirroring the bradykinesia observed in human PD. Furthermore, the presence of rest tremor (Video S3), similar to that reported in PD patients, provides additional evidence of the tree shrew model’s ability to faithfully replicate key PD clinical symptoms, as found in macaque models.

Upon completion of PD modeling in the six tree shrews, we proceeded to quantify their PD motor symptoms. As there are no established scales specifically designed for assessing PD motor symptoms in tree shrews, we utilized the improved *Kurlan* scale (Smith et al., 1993), which is commonly used to assess symptoms in macaque PD research (Imbert et al., 2000). Although this scale was originally developed for Old World monkeys, we deemed it suitable for evaluating PD motor symptoms in tree shrews due to their evolutionary proximity to primates, their complex and flexible movement patterns, and the substantial similarity of symptoms between tree shrew and macaque PD models.

The improved *Kurlan* scale consists of four distinct parts, with part A specifically designed for the assessment of motor symptoms associated with parkinsonism in macaques. It encompasses a comprehensive set of seven scoring items that effectively capture all core motor symptoms of human PD, including bradykinesia, rest tremor, and postural instability, while also considering the unique characteristics of monkeys, such as increased use of upper limbs and defensive responses. Each item is assigned a score, resulting in a total score range of 0 to 20, with higher scores indicating more severe PD symptoms. Preliminary analysis revealed that all seven scoring items were clearly observable in the tree shrews, thereby establishing the feasibly of using the improved *Kurlan* scale to quantitatively evaluate PD motor symptoms in tree shrews.

Quantitative analysis revealed that the PD scores of the six tree shrews rapidly increased to 8–10 at postoperative week 2. From weeks 2 to 20, the PD scores of each tree shrew exhibited a consistent trend with no significant change (Figure 4A). The overall average PD scores of the six tree shrews also exhibited a stable trend with no significant change during the modeling process (Figure 4B). These findings indicate that the six tree shrews displayed continuous and stable PD motor symptoms for approximately five months, highlighting the great potential of this model for future research applications.

**Figure 4:**
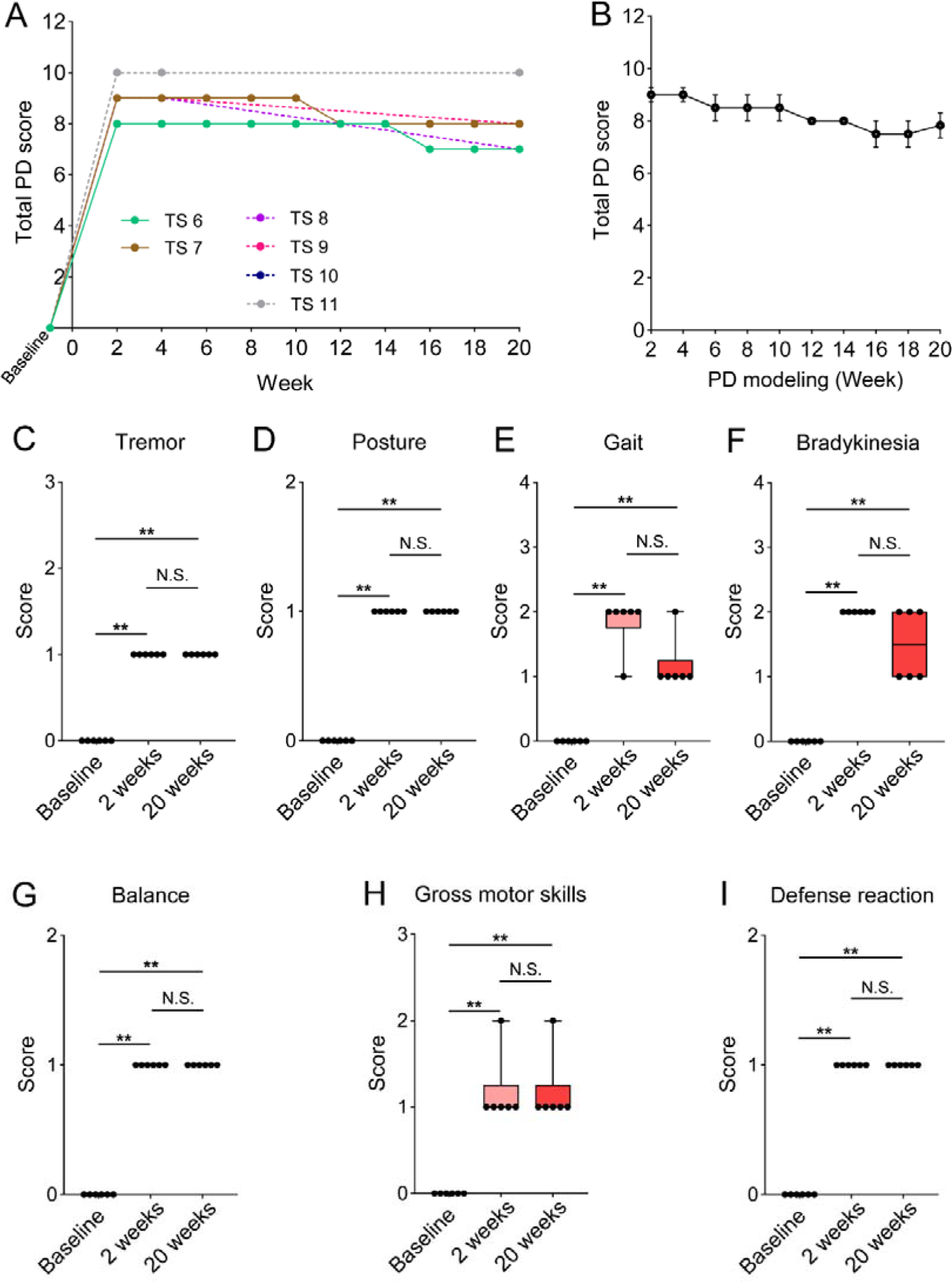
Improved *Kurlan* scale-quantification of clinical motor symptoms of PD in tree shrews following single-dose injection of MPP^+^. (A) Trends in PD scores over time in six tree shrews (TS 6–TS 11) following 50-μg dose MPP^+^ injection into the unilateral substantia nigra. At two weeks post operation, the tree shrews showed typical PD motor symptoms (PD scores 8–10), with no significant change in PD scores for the following five months. (B) Average total PD scores from the six tree shrews showed a consistent trend over time, with no significant differences during modeling (*P* = 0.11). Values are mean ± standard error of the mean (SEM). (C-I) Average scores of the seven individual items comprising the total PD score were compared among the six PD tree shrews at baseline and weeks 2 and 20 of PD modeling. Tremor, abnormal posture, gait, bradykinesia, balance, gross motor skill, and defense reaction scores were significantly higher in tree shrews at weeks 2 and 20 than at baseline (** *P* < 0.01), indicating that the tree shrews developed all PD motor symptoms defined by the improved *Kurlan* scale. Furthermore, there were no significant differences in the scores between weeks 2 and 20, suggesting that the individual PD motor symptoms remained stable for at least five months in all tree shrews. Boxplots (C-I) depict median, 25^th^, and 75^th^ quartiles, and whiskers depict full ranges.

Following the modeling period, the individual scores contributing to the overall PD score of the tree shrews at baseline and weeks 2 and 20 were compared. Notably, no significant differences were observed in any of the scores, including tremor, posture, gait, bradykinesia, balance, gross motor skills (primarily related to upper limb difficulties), and defense reaction, between weeks 2 and 20 of PD modeling, although all scores were significantly elevated compared to baseline (Figure 4C-I). These findings demonstrate that the motor symptoms of PD in tree shrews can be effectively quantified using the classical PD scale for Old World monkeys. Furthermore, our results indicate that the tree shrew PD model more closely resembles that of monkeys than the rodent model, as PD rodents fail to exhibit any of the core clinical symptoms of human PD.

### APO-induced classic rotations in tree shrews injected with MPP^+^

The PD score described above allows for direct evaluation of core clinical motor symptoms in the tree shrew PD model. Rotation induced by APO, an agonist of the dopamine D1 receptor, is a classical pharmacological test for assessing dopaminergic neuronal loss and dysfunction in the substantia nigra pars compacta in animal models of PD with unilateral nigral lesions (Konieczny et al., 2017).

In typical PD animal models with over 90% loss of dopaminergic neurons on the lesioned side, the administration of APO usually results in significant rotational behavior. The direction of rotation can vary depending on factors such as neurotoxin type, dose, and injection site (Carriere et al., 2016; Da Cunha et al., 2008; Dombrowski et al., 2010; Sindhu et al., 2006). In this study, two weeks after surgery, all six tree shrews (TS 6–TS 11) demonstrated APO-induced rotation, indicating substantial loss of dopaminergic neurons (>90%) in the substantia nigra on the lesioned side. Figure 5 and Video S5 show the rotational behavior of one of the six tree shrews (TS 11). APO was administered via hindlimb intramuscular injection (1 mg/mL, 0.2 mL).

**Figure 5:**
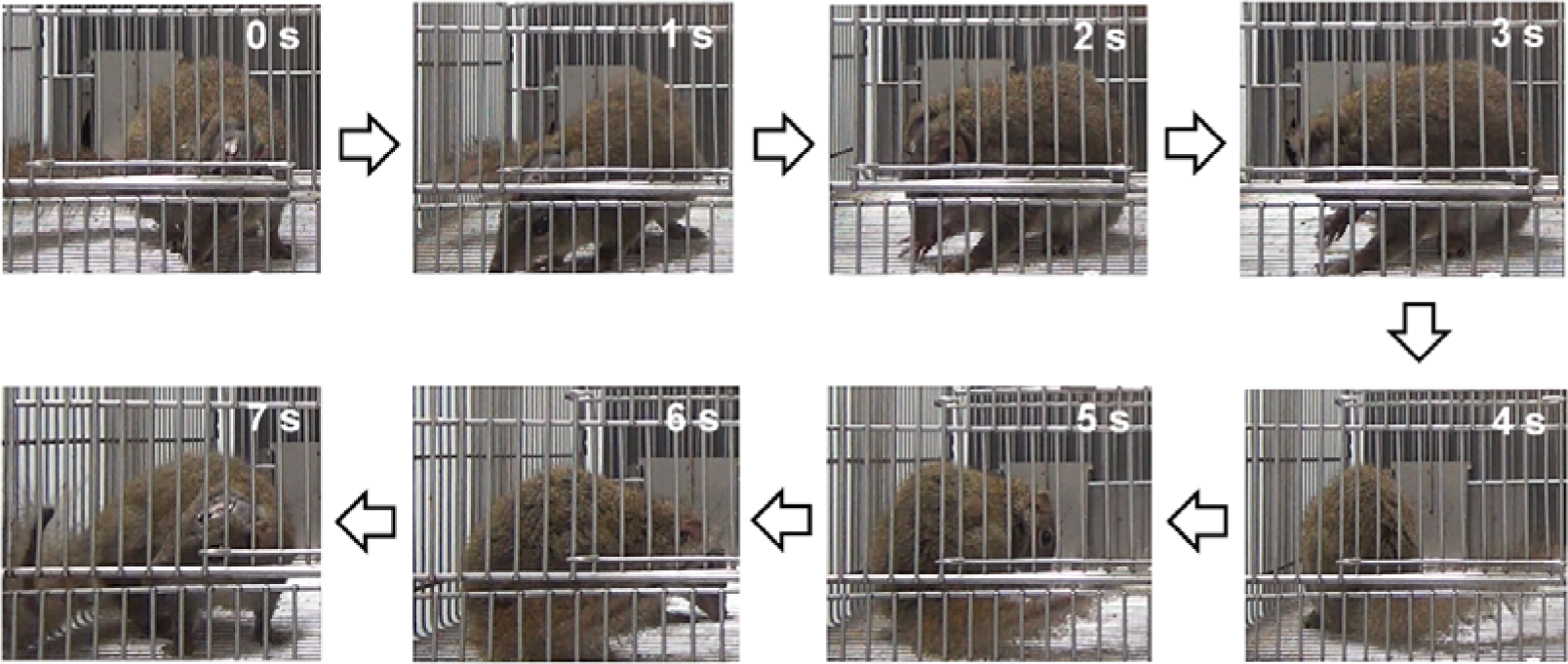
Continuous video recording of APO-induced rotation behavior in TS 11. Rotation lasted for 7 s, during which time TS 11 rotated 360° to its right side. Body position at the start of rotation was defined as 0°. From 1 to 3 s, TS 11 rotated 90° to its right side. From 4 to 5 s, TS 11 rotated 180° to its right side. Finally, from 6 to 7 s, TS 11 continued to rotate 180° to its right side, completing a full 360° turn to its right side.

Figure 6A presents a quantitative analysis of the rotational behavior exhibited by the six tree shrews. Significant increases in APO-induced rotations were observed in all six tree shrews at weeks 2 and 20 after surgery compared to baseline (** *P* < 0.01). These findings provide evidence of more than 90% loss of dopaminergic neurons in the substantia nigra on the lesioned side of the tree shrews. At week 20 of PD modeling, the tree shrews still exhibited stable rotational behavior following intramuscular APO administration, indicating that the damage to dopaminergic neurons in the right substantia nigra could not be restored, as there was no significant difference in the number of APO-induced rotations between weeks 2 and 20 of PD modeling (*P* > 0.05). Figure 6B shows the sustained APO-induced rotations of TS 6 and TS 7 across multiple sampling points from weeks 2 to 20 of PD modeling, with consistent trends observed.

**Figure 6:**
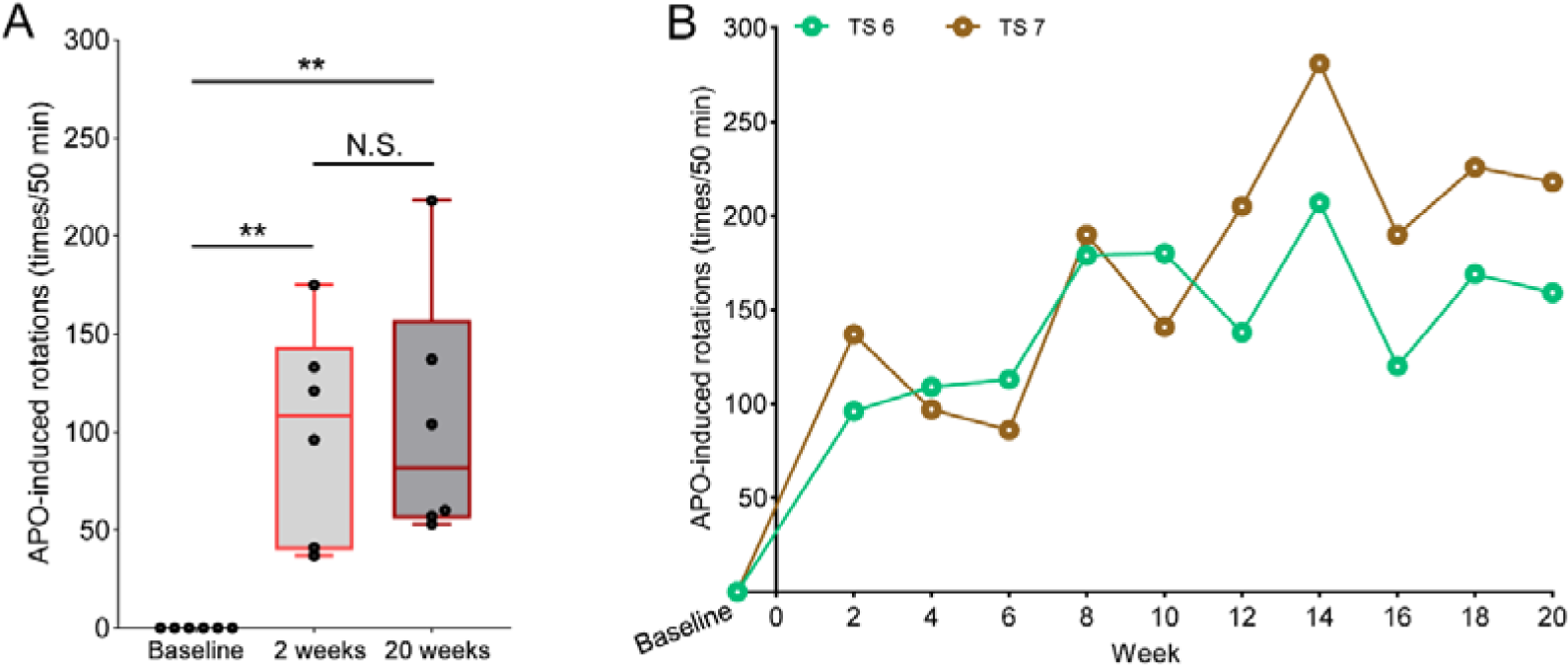
Quantitative analysis of APO-induced rotational behavior in tree shrews following MPP^+^ injection into the unilateral substantia nigra. (A) Two weeks after MPP^+^ was injected into the lateral substantia nigra to model hemi-parkinsonism, all six tree shrews exhibited APO-induced rotations, with significantly higher rotation counts than at baseline (** *P* < 0.01). Prior to modeling, none of the tree shrews exhibited rotations. APO-induced rotation persisted for up to 20 weeks, with the number of rotations significantly higher at week 20 than at baseline (** *P* < 0.01). Number of rotations did not differ significantly (*P* > 0.05) between weeks 2 and 20, indicating that the APO-induced rotations were stable and long-lasting. Boxplots in Panel (A) display median, 25^th^, and 75^th^ quartiles, and whiskers depict full ranges. (B) Rotational behavior of TS 6 and TS 7 at each two-week interval from weeks 2 to 20 after modeling. Both animals exhibited consistent APO-induced rotations at each testing point. Unfortunately, data for TS 8–TS 11 were not sampled due to the COVID-19 pandemic.

In summary, our integrated behavioral observations, clinical PD symptom evaluations, and pharmacological validations demonstrated that the tree shrew model of hemi-parkinsonism, induced through MPP^+^ lesions in the unilateral substantia nigra, consistently manifested all core motor symptoms of PD. Notably, the tree shrew PD model exhibited motor symptoms very similar to those of the macaque PD model for over five months, including bradykinesia, rest tremor, and posture instability.

### Tree shrews exhibited marked nigral dopaminergic neuronal loss, the key pathological feature of PD

Upon completion of the behavioral tests, we characterized the pathological features of the MPP^+^-injected tree shrews. Morphological alterations and significant loss of dopaminergic neurons in the substantia nigra pars compacta are critical pathological features of PD patients (Poewe et al., 2017). In the primate brain, the substantia nigra pars compacta exhibits an abundance of dopaminergic neurons, characterized by a high concentration of TH in the cytoplasm. Consequently, TH immunohistochemical staining serves as a classical method for visualizing and identifying the morphology and number of dopaminergic neurons in the primate substantia nigra (Nagatsu et al., 2019). Here, the tree shrew brains were harvested at week 20 after modeling and processed as coronal substantia nigra sections for TH immunohistochemical staining.

TH immunostaining, with hematoxylin-counterstained nuclei, revealed the pathological features of the dopaminergic neurons in the substantia nigra of the tree shrews. The cell bodies and neurites of the dopaminergic neurons appeared brown, while the nuclei appeared blue, confirming that conventional TH immunohistochemical methods are compatible with tree shrews (Figure 7A). Furthermore, there was almost complete dopaminergic neuronal loss in the substantia nigra on the MPP^+^-lesioned side of the tree shrews (Figure 7B). In contrast, the structure of the substantia nigra pars compacta on the healthy side (saline injected) was intact, with a well-defined substantia nigra pars compacta (SNC, Figure 7C, black dotted area) and substantia nigra pars reticulata (SNR, Figure 7C, light blue dotted area).

**Figure 7:**
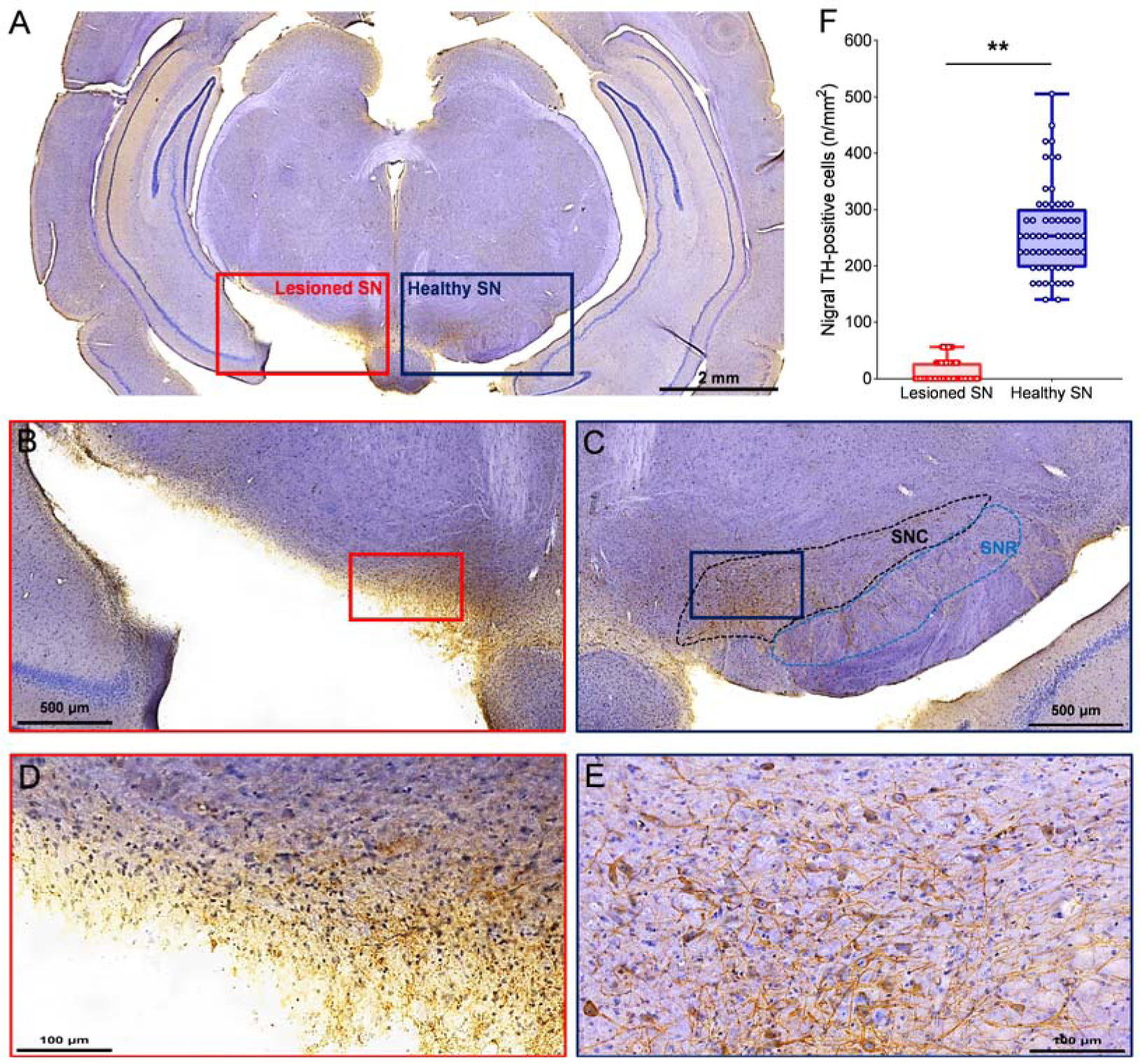
Key pathological features of PD in MPP^+^-lesioned tree shrews. (A) Coronal brain section from TS 10 with bilateral TH immunohistochemical staining of the substantia nigra, with hematoxylin counterstaining of the nuclei. Soma and processes of dopaminergic neurons in the substantia nigra appear brown and nuclei appear blue. (B) Magnified image of red boxed area in Figure A of the substantia nigra on the MPP^+^-lesioned side, showing indistinguishable anatomical morphology. (C) Magnified image of blue boxed area in Figure A of the substantia nigra on the healthy side, showing structurally intact substantia nigra pars compacta (SNC, black dotted area) and pars reticulata (SNR, light blue dotted area). (D) Magnified image of red boxed area in Figure B on the MPP^+^-lesioned side, showing total loss of dopaminergic neurons in the substantia nigra. (E) Magnified image of blue boxed area in Figure C on the healthy side, showing many dopaminergic neurons with intact somata, with well-developed TH-positive brown-stained processes and clear blue-stained nuclei. Images (B) and (C), (D), and (E) were symmetrical in anatomical location. (F) In total, 60 midbrain coronal sections from six tree shrews containing bilateral areas of the substantia nigra showed that dopaminergic neuronal density was significantly lower on the MPP^+^-lesioned side than on the healthy side (** *P* < 0.01), with nearly 95% loss. Boxplots (F) depict median, 25^th^, and 75^th^ quartiles, and whiskers depict full ranges.

Upon examination under 20× magnification, only a few surviving dopaminergic neurons were observed in the MPP^+^-lesioned side of the substantia nigra (Figure 7D). In contrast, many well-defined and TH-positive dopaminergic neurons were present in the healthy side of the substantia nigra. The somas of these neurons were mostly spindle-shaped, with clearly visible brown-stained processes and blue-centered nuclei (Figure 7E), similar to the recently published study on midbrain anatomy and cellular morphology in normal tree shrews (Ni et al., 2021).

We observed macroscopically visible lesions in the substantia nigra on the lesioned side (sampled anterior to posterior) of the brains of all six tree shrews. At the microscopic level, we observed an almost complete loss of nigral cells. These observations confirmed the precision of MPP^+^-induced lesions to the substantia nigra and demonstrated that the tree shrew MPP^+^ lesion model can replicate the core pathological hallmark of substantial nigral dopaminergic neuronal loss, a fundamental characteristic of human PD. To calculate the density of dopaminergic neurons in the substantia nigra, we counted the number of TH-positive neurons per unit area (Figure 7DE) in symmetric regions between lesioned and healthy substantia nigra (Figure 7BC). Under 40× magnification, the density of TH-positive neurons in the substantia nigra on the MPP^+^-lesioned side was significantly lower than that on the healthy side (** *P* < 0.01), with nearly 95% loss (Figure 7F). Moreover, the concentration of dopaminergic neurons in the substantia nigra on the non-lesioned side of the tree shrews was analogous to that observed in macaques (Chu et al., 2019; Li et al., 2021a; Li et al., 2021b).

In summary, unilateral MPP^+^ lesions in tree shrews successfully recapitulates the key pathological features of human PD, including marked morphological modifications and severely depleted dopaminergic neurons in the substantia nigra.

### Size comparison of nigral dopaminergic neurons reveals greater similarity between tree shrews and macaques than rodents

Based on the observed PD behaviors and pathological findings, our tree shrew PD model, established with unilateral substantia nigra MPP^+^-induced lesions, closely resembled the hemi-parkinsonian macaque model, well replicating the core phenotype of human PD and outperforming rodent PD models in terms of clinical symptoms and pathological hallmarks. As such, we explored why the tree shrew, as a potential alternative to nonhuman primates, could be advantageous for PD modeling. Following the principle that structure determines function, we hypothesized that the contrasting morphology of nigral dopaminergic neurons across these species may play a contributing factor. Thus, we conducted a systematic comparison of the morphological features of nigral dopaminergic neurons in tree shrews, C57 mice, Wistar rats, and macaques to identify similarities and differences.

Referring to previously published morphological data on dopaminergic neurons in the substantia nigra of C57 mice (Gonzalez-Cabrera et al., 2017; Lin et al., 2020; Tiklova et al., 2019; Won et al., 1989) and Wistar rats (Bigham et al., 2021; De March et al., 2006; Guan et al., 2000; Karunasinghe et al., 2017; Van der Perren et al., 2011), we determined that the average soma diameter of dopaminergic neurons in the substantia nigra of the C57 mice was 16.5 μm, very close to the reported value (16.4 μm) (Nelson et al., 1996), while the average diameter in the Wistar rats was 17.3 μm, also close to the previously reported value (16.5 μm) (German and Manaye, 1993). We used the same measurement to compare morphological parameters, including mean diameter of the soma and nucleus, of dopaminergic neurons in the substantia nigra of C57 mice, Wistar rats, six tree shrews, and six macaques (used in our other experiments).

Average diameters of the soma and nucleus of dopaminergic neurons in the substantia nigra of tree shrews were significantly greater than those observed in the C57 mice and Wistar rats, but more comparable to those seen in macaques (Figure 8AC). To facilitate size comparison of dopaminergic neurons in the substantia nigra among the species, we converted the size parameters of C57 mice, Wistar rats, and tree shrews into percentages relative to those of macaques. Results showed that nigral dopaminergic neurons were larger in tree shrews than in C57 mice and Wistar rats, and closer in scale to those observed in macaques. Specifically, average soma diameter in the tree shrew was 89.33% of that observed in macaques, but only 60.94% and 64.01% of that observed in mice and rats (Figure 8B).

**Figure 8:**
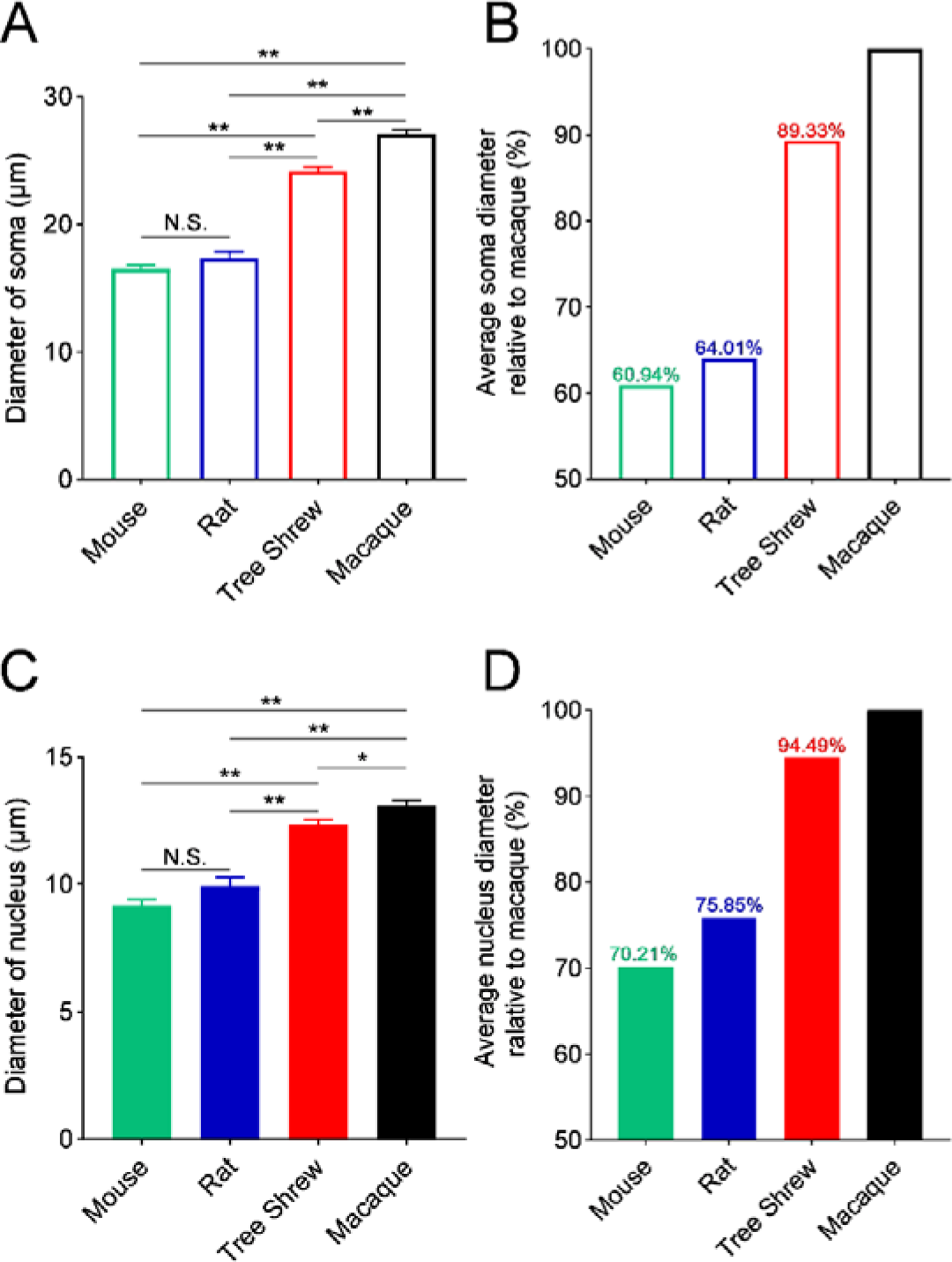
Comparison of morphological parameters of nigral dopaminergic neurons in C57 mice, Wistar rats, tree shrews, and macaques. (A) Average soma diameter of nigral dopaminergic neurons in mice, rats, tree shrews, and macaques, with diameter in tree shrews being much larger than that of rodents, but relatively close to that of macaques. (B) Conversion of average soma diameter of C57 mice, Wistar rats, and tree shrews (numerators) relative to macaques (denominator) (as a percentage). Results revealed that the cell body scale of nigral dopaminergic neurons in tree shrews was closer to that in macaques than in rodents. (C) Average nuclear diameter of the nigral dopaminergic neurons in mice, rats, tree shrews, and macaques. Results showed that diameter in tree shrews was much larger than that in rodents and relatively close to that in macaques. (D) Size parameters of C57 mice, Wistar rats, and tree shrews relative to macaques (as a percentage). Results revealed that the nucleus size of nigral dopaminergic neurons in tree shrews is closer to that in macaques than in rodents. Values in A and C are presented as mean ± SEM.

Our analysis of dopaminergic neurons revealed that the nuclear diameter in tree shrews was 94.49% of that observed in macaques. In contrast, the C57 mice and Wistar rats exhibited much smaller nuclear diameters (70.21% and 75.85% of that observed in macaques) (Figure 8D). These findings highlight the morphological similarity in the size of nigral dopaminergic neurons between tree shrews and macaques, distinguishing them from rodents. Notably, this difference in size is not attributable to differences in body size, as tree shrews and Wistar rats share similar size and weight characteristics, yet the nigral dopaminergic neurons in tree shrews were significantly larger than those in rats.

Based on the principle that structure plays a significant role in determining function, we hypothesize that the similarity in dopaminergic neuronal morphology between tree shrews and macaques may serve as a biological basis for the expression of core phenotypes in tree shrews that closely resemble those observed in monkey PD models.

## Discussion

In this study, we have successfully developed a PD model in tree shrews by precisely injecting MPP^+^ into the right nigral region, resulting in specific and effective damage to more than 95% of local dopaminergic neurons (Figure 7). Using the improved *Kurlan* scale for the assessment of clinical motor symptoms in PD macaques, we showed that the tree shrew model exhibited and maintained all core clinical symptoms of PD closely resembling those observed in macaques over a period of five months, including bradykinesia, rest tremor, and postural instability (Figure 4). The resemblance between the two models is striking to the extent that distinguishing the motor symptoms of PD in tree shrews from those in macaques based solely on the PD scores was challenging, despite our extensive experience in creating drug-induced PD models in macaques. This indicates that the model well replicates the core clinical symptoms of human PD. In addition, the tree shrew also exhibited and maintained APO-induced rotation, a classical marker of the hemi-parkinsonian model (Figures 5–6).

We believe that the utilization of the tree shrew for PD modeling offers a significant advantage in terms of its movement dimension and flexibility. Similar to macaques, tree shrews naturally inhabit a much wider range of three-dimensional space. Moreover, tree shrews are extremely mobile, and their movements exhibit a level of flexibility and complexity that is comparable to that of macaques. This inherent biological characteristic provides the basis for the tree shrew PD model to accurately replicate the symptoms observed in macaque PD models. In contrast, due to their distinct evolutionary background, rodents tend to exhibit daily behaviors that are predominantly restricted to a two-dimensional plane, with markedly less complex and flexible movement in comparison to macaques. As such, the commonly used rodent PD models do not possess classical PD motor symptoms, particularly rest tremor. Motor impairments in rodents are also difficult to quantify in a scale-scored manner (Beal, 2001), and often require indirect behavioral paradigms such as the rotarod task and pole test (Cunha et al., 2017). Compared to rodents, tree shrews exhibit distinct advantages in mimicking the core clinical symptoms of PD patients and have the potential to partially replace macaque PD models.

Based on analysis of cell morphology, we found that dopaminergic neurons in the substantia nigra of tree shrews closely resemble those of macaques, particularly in comparison to rodents (Figure 8). Specifically, average soma and nucleus diameter in tree shrews was 89.33% and 94.49% of that in macaques, respectively, while in rodents, these measures were less than 80% of that in macaques. Considering the established principle that structure plays a crucial role in determining function, this morphological similarity between tree shrews and macaques may serve as a structural foundation for the superior ability of the tree shrew PD model to faithfully replicate the core clinical symptoms of PD when compared to rodent models.

The tree shrew model of PD also benefits from its distinctive modeling approach. The direct injection of MPP^+^ into the unilateral substantia nigra enables straightforward and efficient establishment of the model, with a short modeling period and high consistency in nigral lesion creation. As a result, the model effectively exhibits all classical PD motor symptoms and maintains their stability over time.

The limited availability and high costs associated with nonhuman primate resources pose challenges for their widespread use in research. In contrast, tree shrews offer distinct advantages, such as ease of breeding and shorter growth cycles (Yao, 2017). Therefore, the use of our tree shrew model of PD offers the opportunity to conduct large-scale experiments that can provide an initial evaluation of the efficacy and safety of novel treatments for PD. Additionally, this model has the potential to generate valuable prospective data that can facilitate the advancement of clinical translation in the field of PD research.

In conclusion, our tree shrew PD model is a promising alternative to the macaque PD model, offering generalizability and the potential for partial replacement. Furthermore, this model provides a novel platform for evaluating the efficacy of new therapeutic strategies and for studying the neural mechanisms underlying PD.

## Author Contributions

Xintian Hu, Lin Xu, Jiali Li, Bingyin Shi and Christoph W. Turck conceived the idea and designed the experiments. Hao Li, Leyi Mei, Xiupeng Nie, and Lingping Wu conducted the experiments. Longbao Lv, Jitong Yang, Haonan Cao, Jing Wu, Yuhua Zhang, Yingzhou Hu, and Wenchao Wang collected and analyzed the data. Hao Li, Leyi Mei, Xiaofeng Ren, Christoph W. Turck, and Xintian Hu drafted and critically revised the manuscript. All authors approved the final version of the manuscript to be published and agreed to be accountable for all aspects of the work.

## Acknowledgments

We would like to thank technician Hongwei Li for help in data collection. We also thank the staff from tree shrew breeding team of the Kunming Institute of Zoology, Chinese Academy of Sciences, for their kind help in animal care. This study was partially supported by the National Key Research and Development Program of China (2021YFF0702700, 2018YFA0801403), STI2030-Major Projects (2021ZD0200900), Key-Area Research and Development Program of Guangdong Province (2019B030335001), National Natural Science Foundation of China (81941014, 81960422), Strategic Priority Research Program of the Chinese Academy of Sciences (XDB32060200), STI2030-Major Projects (2022ZD0205200, 2022ZD0212700), Yunnan Fundamental Research Projects (202201AT070153, 202201AT070139, 202001AT070130), Science and Technology Project of Yunnan Province (202101AY070001-001), CAS “Light of West China” Program, and “Yunnan Revitalization Talents Support Plan”.

